# Tethering Piezo channels to the actin cytoskeleton for mechanogating via the E-cadherin-β-catenin mechanotransduction complex

**DOI:** 10.1101/2020.05.12.092148

**Authors:** Jing Wang, Jinghui Jiang, Xuzhong Yang, Li Wang, Bailong Xiao

## Abstract

The mechanically activated Piezo channel plays a versatile role in conferring mechanosensitivity to various cell types. However, how it incorporates its intrinsic mechanosensitivity and cellular components to effectively sense long-range mechanical perturbation across a cell remains elusive. Here we show that Piezo1 is biochemically and functionally tethered to the actin cytoskeleton via the E-cadherin-β-catenin mechanotransduction complex, whose perturbation significantly impairs Piezo1-mediated responses. Mechanistically, the adhesive extracellular domain of E-cadherin interacts with the cap domain of Piezo1 that controls the transmembrane gate, while its cytosolic tail might interact with the cytosolic domains of Piezo1 that are in close proximity to its intracellular gates, allowing a direct focus of adhesion-cytoskeleton-transmitted force for gating. Specific disruption of the intermolecular interactions prevents cytoskeleton-dependent gating of Piezo1. Thus, we propose a force-from-filament model to complement the previously suggested force-from-lipids model for mechanogating of Piezo channels, enabling them to serve as versatile and tunable mechanotransducers.

**Highlights:**

- Revealed biochemical and functional interactions between Piezo1 and the E-cadherin-β-catenin-F-actin mechanotransduction complex.
- Identified critical mechanogating domains of Piezo1 as E-cadherin binding domains.
- Specific disruption of the intermolecular interactions between Piezo1 and E-cadherin prevents cytoskeleton-dependent gating of Piezo1.
- Proposed a tether model for mechanogating of Piezo channels.

## Introduction

The cellular mechanotransduction process requires designated mechanotransducers to convert mechanical force into various fundamental biological activities, ranging from cellular division, differentiation and migration to physiological sensing of blood pressure, touch, mechanical pain, balance and sound (Chalfie, 2009; Douguet and Honore, 2019; Jin et al., 2020; Katta et al., 2015; Leckband and de Rooij, 2014; Lecuit and Yap, 2015; Niessen et al., 2011). Adhesion molecules and mechanosensitive ion channels represent two major types of mechanotransducers. Adhesion-based mechanotransduction utilizes adhesion molecules such as E-cadherin, a member of the type I classical cadherin family that contains an ectodomain composed of five tandem extracellular cadherin (EC) repeats involved in Ca^2+^ dependent homotypic interactions, a single-pass transmembrane region (TM), and a conserved cytoplasmic region with binding sites for β-catenin, to physically link to the actin cytoskeleton via actin-binding proteins such as vinculin (Leckband and de Rooij, 2014; Lecuit and Yap, 2015; Niessen et al., 2011). Such an organization allows reciprocal force transmission and signal transduction between cadherin and cytoskeleton, which is critical for maintaining the integrity of many multicellular tissues (Leckband and de Rooij, 2014; Lecuit and Yap, 2015; Niessen et al., 2011).

Mechanosensitive ion channels can respond rapidly to mechanical forces to allow ion flux via two main proposed mechanogating models: force-from-lipids and force-from-filament (Douguet and Honore, 2019; Gillespie and Walker, 2001; Jin et al., 2020; Kung et al., 2010; Markin and Hudspeth, 1995; Ranade et al., 2015). The force-from-lipids model proposes that the channel can directly respond to changes in membrane tension (Anishkin et al., 2014; Kung et al., 2010), which is exemplified by the bacteria mechanosensitive channel of large conductance (MscL) (Martinac et al., 1990; Sukharev et al., 1994) and the eukaryotic two-pore domain K^+^ channels TREK-1 and TRAAK (Brohawn et al., 2014). In contrast, the force-from-filament model or tether model proposes that the channel is physically tethered to either extracellular matrix or intracellular accessory structures such as cytoskeleton for mechanogating (Gillespie and Walker, 2001; Markin and Hudspeth, 1995). The fly NOMPC channel has been elegantly demonstrated to interact with microtubules for a tethered gating model (Zhang et al., 2015). The tether model also raises an intriguing mechanism for integrating adhesion-based and channel-based mechanotransduction systems via converging on cytoskeleton. However, no mammalian mechanosensitive ion channel has yet to be clearly demonstrated to adopt such a tether model.

The mechanically activated Piezo channel family, including Piezo1 and Piezo2 in mammals (Coste et al., 2010; Coste et al., 2012), represents a bona fide class of eukaryotic mechanotransduction cation channels (Murthy et al., 2017; Wu et al., 2016; Xiao, 2019). Heterologously expressed Piezo1 is characteristically activated by various forms of mechanical stimulation, including poking, stretching, shear stress, and substrate deflection (Bavi et al., 2019; Coste et al., 2010; Cox et al., 2016; Lewis and Grandl, 2015; Li et al., 2014; Poole et al., 2014). Cell-attached patch-clamp recordings from Piezo1-transfected HEK293T cells have measured a T_50_ (the tension required for half maximal activation) of ~1.4 mN/m for activation of Piezo1 by lateral membrane tension (Lewis and Grandl, 2015), demonstrating its extraordinary mechanosensitivity. Endogenously expressed Piezo1 or Piezo2 confers exquisite mechanosensitivity to various cell types (Murthy et al., 2017; Xiao, 2019), including epithelial cells (Eisenhoffer et al., 2012; Gudipaty et al., 2017), endothelial cells (Li et al., 2014; Nonomura et al., 2018; Ranade et al., 2014a), smooth muscle cells (Retailleau et al., 2015), red blood cells (Zarychanski et al., 2012), chondrocytes (Servin-Vences et al., 2017), osteoblasts (Sun et al., 2019), merkel cells (Woo et al., 2014) and sensory neurons (Ranade et al., 2014b). Such Piezo-mediated mechanotransduction process controls a wide variety of key biological activities (Douguet and Honore, 2019; Murthy et al., 2017; Xiao, 2019), including epithelial homeostasis (Eisenhoffer et al., 2012; Gudipaty et al., 2017), vascular and lymphatic development and remodeling (Li et al., 2014; Nonomura et al., 2018; Ranade et al., 2014a), blood pressure regulation (Rode et al., 2017; Wang et al., 2016; Zeng et al., 2018), bone formation (Sun et al., 2019), and somatosensation of touch (Ranade et al., 2014b; Woo et al., 2014), balance (Woo et al., 2015), tactile pain (Murthy et al., 2018; Szczot et al., 2018) and breathing (Nonomura et al., 2017). Abnormal Piezo channel functions arising from genetic mutations have been associated with many human genetic diseases, including *Piezo1*-based dehydrated hereditary stomatocytosis (Zarychanski et al., 2012) and familial generalized lymphatic dysplasia (Fotiou et al., 2015; Lukacs et al., 2015), and *Piezo2*-based distal arthrogryposis (Coste et al., 2013), scoliosis and peripheral sensory dysfunction such as loss of tactile pain (Chesler et al., 2016; Szczot et al., 2018). Given their prominent roles in cellular and physiological mechanotransduction and therapeutic potential as validated drug targets (Xiao, 2019), it is pivotal to understand how Piezo channels function as versatile mechanotransducers for effectively sensing and converting distinct forms of mechanical stimuli into electrochemical signals under variable cellular contexts.

Mouse Piezo1 and Piezo2 are large membrane proteins of over 2500 amino acids, and adopt a unique 38-transmembrane-helix (TM) topology, in which the first 36 TMs folded into 9 structurally repetitive transmembrane helical units (THU) of 4-TMs each and the last two TMs forming the ion-conducting pore module (Wang et al., 2019; Zhao et al., 2018a). The homotrimeric Piezo1 or Piezo2 channel resembles a gigantic three-bladed propeller-like structure (Ge et al., 2015; Guo and MacKinnon, 2017; Saotome et al., 2018; Wang et al., 2019; Zhao et al., 2018a), in which the non-planar transmembrane regions of 114 TMs in total are collectively curved into a unique nano-bowl shape with a mid-plane bowl surface area of 700 nm^2^ and a projected in-plane are of 450 nm^2^ (Wang et al., 2019), which might cause the residing membrane to adopt a similar highly curved and non-planar shape (Guo and MacKinnon, 2017; Lin et al., 2019) (Fig. 1A). The unusual nano-bowl configuration of the Piezo protein-membrane system has led to the intriguing hypothesis that tension-induced flattening of the mechanosensing-blades and concurrent expansion of the in-plane membrane area might provide sufficient gating energy to gate the ion-conducting pore (Guo and MacKinnon, 2017; Haselwandter and MacKinnon, 2018; Lin et al., 2019; Saotome et al., 2018; Zhao et al., 2018a). Such an intrinsic Piezo structure-based mechanism might confer an exquisite mechanosensitivity to the Piezo channel in response to changes in local membrane curvature and tension. When reconstituted into a two-droplet bilayer system, Piezo1 was activated in an asymmetric bilayer but not in a symmetric bilayer (Syeda et al., 2016), indicating its intrinsic sensitivity to membrane curvature. Furthermore, negative pressured-induced Piezo1 activation has been recorded from HEK293 cell bleb membranes (Cox et al., 2016). On the basis of the signature bowl-shaped feature of the Piezo-membrane system and the electrophysiological characterizations, the Piezo channel might adopt a force-from-lipids gating mechanism to enable a versatile response to changes in local curvature and membrane tension.

**Figure 1.**
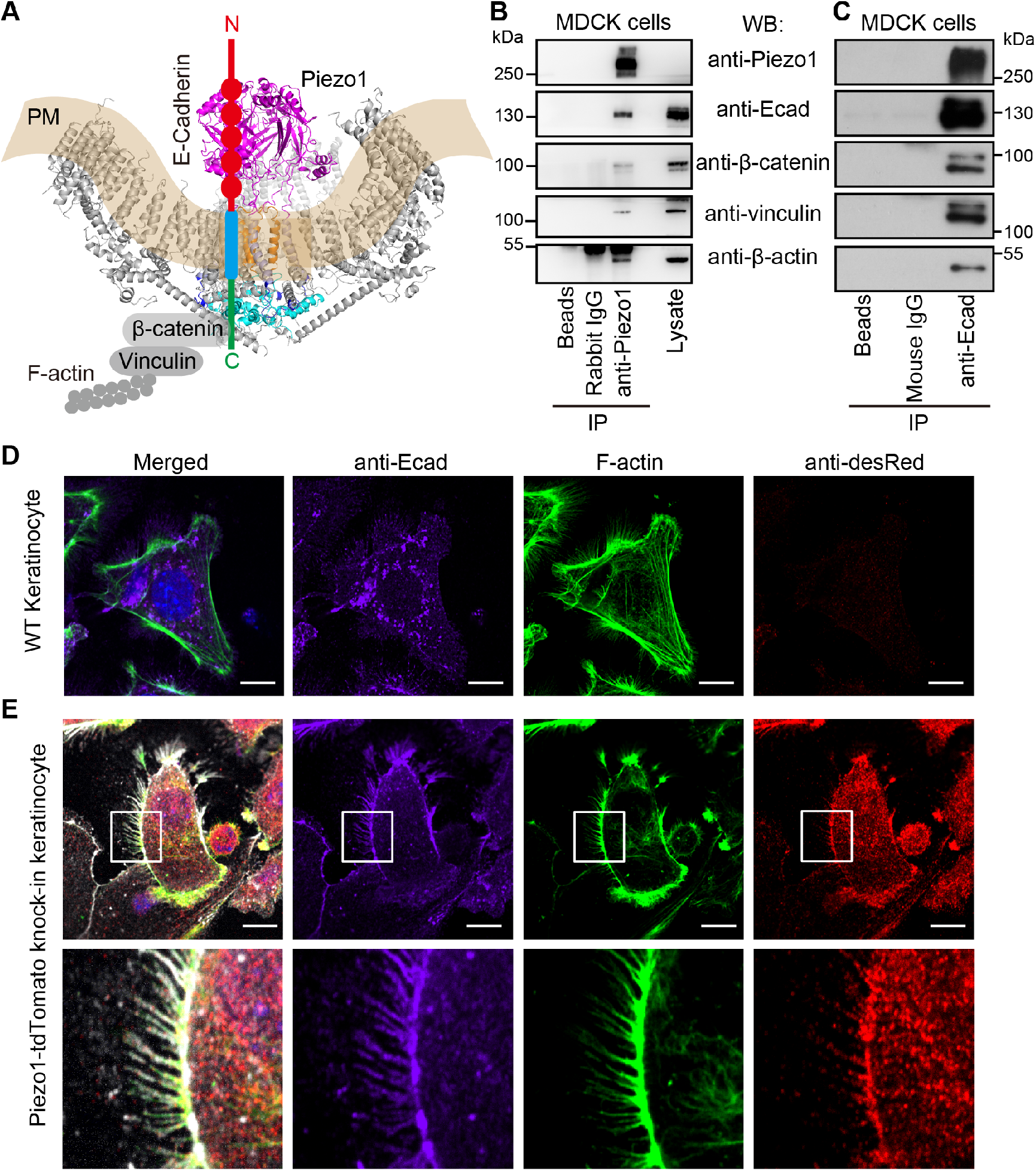
Endogenous interaction of Piezo1 with the E-cadherin/β-catenin/vinculin/F-actin mechanotransduction complex. (A) Paradigm showing the Piezo1 structure (PDB: 5z10) and the E-cadherin/β-catenin/vinculin/F-actin mechanotransduction complex. The transparent yellow area indicates the nano-bowl-shaped plasma membrane curved by the highly non-planar transmembrane region of the Piezo1 structure. The extracellular cap embedded at the center of the nano-bowl shape of Piezo1 is shown in purple. The N-terminal ectodomain composed of 5 tandem cadherin domains, the single-pass transmembrane region and the C-terminal cytosolic tail of E-cadherin are colored in red, blue and green, respectively. (B, C) Representative western blots showing co-immunoprecipitation of endogenously expressed Piezo1, E-cadherin, β-catenin, β-actin in MDCK cells using either the anti-Piezo1 (B) or anti-E-cadherin antibody (C) for immunoprecipitation. (D, E) Representative immunofluorescent staining images of endogenous expressed Piezo1-tdTomato fusion proteins, E-cadherin and phalloidin-stained F-actin fibers in primarily cultured keratinocytes either derived from wild-type control mice (D) or from the Piezo1-tdTomato knock-in mice (E). The culture medium contains 0.07 mM Ca^2+^. The region marked with white box in the top row of panel E is enlarged in the bottom row for highlighting the co-location of the proteins at the filopodia structures. White area indicates co-localization of Piezo1-tdTomato, E-cadherin and F-actin. The scale bar represents 10 μm.

Despite the well-established importance of membrane tension for many mechanotransduction processes, including mechanogating of ion channels, it has recently been demonstrated that no long-range propagation of membrane tension occurs in intact cell membranes (Shi et al., 2018), which imposes a challenge for Piezo channels to use the force-from-lipids model to sense remote mechanical perturbation across a cell. Experimentally, Piezo1 can mediate both localized and whole-cell mechanical responses regardless whether the mechanical stimuli are either exogenously applied or endogenously originated (Coste et al., 2010; Ellefsen et al., 2019; Pathak et al., 2014; Shi et al., 2018). In light of these paradoxical experimental observations, we have set out to test whether Piezo channels might utilize a complementary force-from-filament gating model to enable long-distance sensing of mechanical perturbation across an intact cell via cytoskeleton, which is effective in propagating mechanical force either exogenously applied or endogenously generated. Toward this, we have found that Piezo channels are biochemically and functionally tethered to the actin cytoskeleton via the mechanotransduction complex comprising E-cadherin, β-catenin and vinculin, establishing the first mammalian mechanosensitive ion channel undertaking a tether model for mechanogating.

## Results

### Endogenous interaction of Piezo1 with the E-cadherin/β-catenin/vinculin/F-actin mechanotransduction complex

To investigate whether endogenously expressed Piezo1 might interact with cytoskeleton, we first carried out co-immunoprecipitation studies by choosing the Madin-Darby canine kidney (MDCK) cells, which utilize both Piezo1 and the E-cadherin-dependent mechanotransduction complex including β-catenin, vinculin and actin for cellular mechanotransduction processes such as cell extrusion, proliferation and migration (Benham-Pyle et al., 2015; Eisenhoffer et al., 2012; Gudipaty et al., 2017). Using the custom generated anti-Piezo1 antibody (Zhang et al., 2017), we succeeded in pulling down endogenous Piezo1 proteins from MDCK cells as revealed by western blotting (Fig. 1B). Importantly, we also detected E-cadherin, β-catenin, vinculin and actin specifically from the anti-Piezo1 immunoprecipitated sample, but not from control samples pulled down with either the beads alone or the beads bound with the rabbit IgG antibody (Fig. 1B). Reciprocally, immunoprecipitation of E-cadherin using the anti-E-cadherin antibody led to co-immunoprecipitation of Piezo1, β-catenin, vinculin and β-actin (Fig. 1C). Together, these data demonstrate that endogenously expressed Piezo1 proteins biochemically interact with the E-cadherin-mediated mechanotransduction components (Fig. 1A), prompting us to ask whether Piezo1 might be tethered to actin fibers via E-cadherin in cells.

Given that the Piezo1 antibody used for immunoprecipitation and western blotting studies in Fig. 1B, C did not have the required specificity and affinity for staining endogenously expressed Piezo1 proteins (Zhang et al., 2017), we carried out co-immunostaining experiments using primarily cultured keratinocytes derived from the Piezo1-tdTomato knock-in (KI) mice, which express the Piezo1-tdTomato fusion protein (Ranade et al., 2014a). Keratinocytes are epithelial cells and express relatively high level of Piezo1 (Coste et al., 2010). Immunostaining using the anti-dsRed antibody for recognizing the tdTomato fluorescent protein clearly detected the expression of the Piezo1-tdTomato fusion proteins in the Piezo1-tdTomato knock-in keratinocytes (Fig. 1E), but not in cells derived from the wild-type control mice (Fig. 1D). Consistent with its expression pattern observed in mouse neural stem/progenitor cells (mNSPCs) or mouse embryonic fibroblasts (MEFs) derived from the Piezo1-tdTomato knock-in mice (Ellefsen et al., 2019), Piezo1-tdTomato was detected in both the cell membrane and inside the cell, but relatively enriched at the leading edge of the cell membrane, including filopodia protrusions (Fig. 1E). The endogenously expressed E-cadherin was immunostained with the anti-E-cadherin antibody, while the actin fibers were stained with phalloidin (Fig. 1D, E). Since the keratinocytes were cultured in medium containing low Ca^2+^ (70 μM), which does not favor the formation of homotypic interaction of E-cadherin at cell-cell junctions (Lewis et al., 1994), E-cadherin was not restricted to cell-cell junctions, instead also located at cell periphery and filopodia protrusions without cellular contacts (Fig. 1D, E). Importantly, Piezo1, E-cadherin and actin fibers were clearly co-localized at cell periphery and filopodia protrusions free of cellular contacts (Fig. 1E). Together, these data suggest that Piezo1 could be tethered to actin fibers via E-cadherin independently of its homotypic interaction at cell-cell junctions.

### Endogenous regulation of Piezo1 responses by the E-cadherin-β-catenin-actin complex

We next investigated whether the Piezo1 channel function is influenced by the E-cadherin-mediated mechanotransduction complex. Although Piezo1 has been proposed to mediate both crowding-dependent cell extrusion and stretching-induced proliferation of MDCK cells, its channel activity has not been functionally assayed (Eisenhoffer et al., 2012; Gudipaty et al., 2017). Using Fura-2 Ca^2+^ imaging, we found that confluent MDCK cells transfected with the scrambled control siRNA showed robust Ca^2+^ increase in response to the application of 5 μM Yoda1, a gating modifier that potentiates the mechanosensitivity of Piezo1 (Syeda et al., 2015) (Fig. 2A, B). Importantly, the Yoda1-induced response was drastically reduced in MDCK cells transfected with siRNA against Piezo1 (Fig. 2A, B), suggesting a Piezo1-dependent Ca^2+^ response. Interestingly, knockdown of either E-cadherin or β-catenin significantly reduced the Yoda1 response (Fig. 2A, B). In line with the functional reduction of the Yoda1-evoked Piezo1 response, quantitative real-time PCR (RT-PCR) verified significant reduction of mRNA expression of Piezo1, E-cadherin and β-catenin (Fig. 2C).

**Figure 2.**
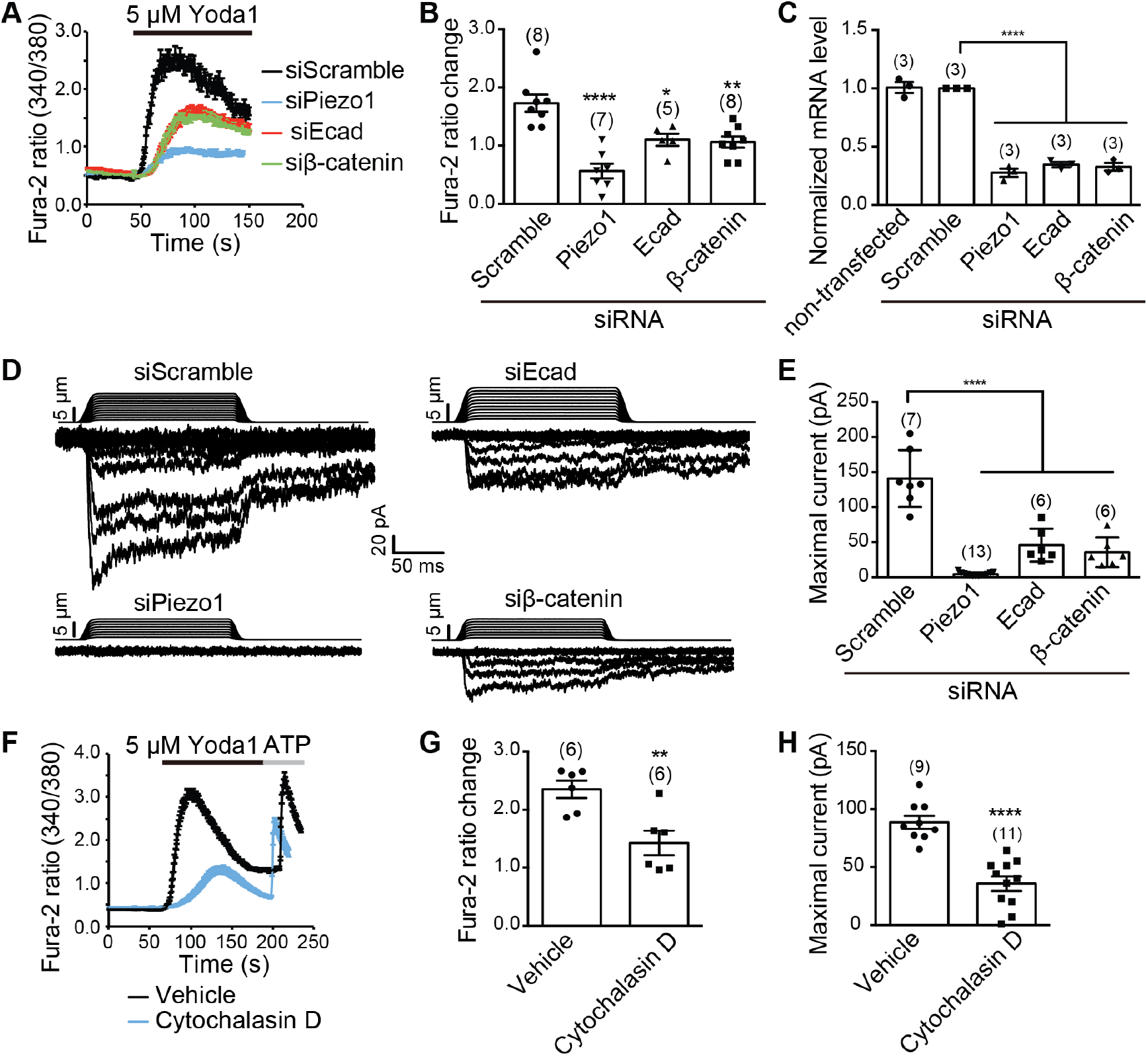
Endogenous regulation of Piezo1 responses by E-cadherin/β-catenin/F-actin. (A) Representative averaged Ca^2+^ imaging traces showing Yoda1-induced change of Fura-2 ratio (340/380) in confluent MDCK cells transfected with the indicated siRNAs. (B) Scatter plot of Yoda1-induced Fura-2 ratio change of MDCK cells transfected with the indicated siRNAs. Each bar represents mean ± s.e.m., and the imaged coverslip number is labeled above the bar. One-way ANOVA with Dunn’s comparison to the group transfected with scramble siRNA. ****P < 0.0001, **P < 0.01, *P < 0.05. (C) Scatter plot of normalized mRNA level of MDCK cells without transfection or transfected with the indicated siRNAs. Each bar represents mean ± s.e.m., and the number of repeats is labeled above the bar. One-way ANOVA with Dunn’s comparison to the group transfected with scramble siRNA. ****P < 0.0001. (D) Representative poking-evoked whole-cell currents of sparsely re-plated MDCK cells transfected with the indicated siRNAs. (E) Scatter plot of the maximal poking-evoked whole-cell currents of MDCK cells transfected with the indicated siRNAs. Each bar represents mean ± s.e.m., and the recorded cell number is labeled above the bar. One-way ANOVA with Dunn’s comparison to the group transfected with scramble siRNA. ****P < 0.0001. (F) Representative averaged single-cell Ca^2+^ imaging traces showing Yoda1-induced change of Fura-2 ratio (340/380) in MDCK cells pre-treated with either vehicle or 10 μM cytochalasin D. (G) Scatter plot of Yoda1-induced Fura-2 ratio change of MDCK cells pre-treated with either vehicle or 10 μM cytochalasin D. Each bar represents mean ± s.e.m., and the imaged coverslip number is labeled above the bar. Unpaired Student’s t-test. **P < 0.01. (H) Scatter plot of the maximal poking-evoked whole-cell currents of MDCK cells pre-treated with either vehicle or 10 μM cytochalasin D. Each bar represents mean ± s.e.m., and the recorded cell number is labeled above the bar. Unpaired Student’s t-test. ****P < 0.0001.

To directly assay Piezo1-mediated mechanically activated whole-cell currents in MDCK cells, we utilized a piezo-driven blunted glass pipette to mechanically probe the cell membrane under a whole-cell recording configuration. To facilitate recording, MDCK cells were re-plated to obtain relatively dispersed cells, which thus were not under a confluent condition. MDCK cells showed slowly adapting and mechanically activated currents with an averaged maximal current of 140.9 ± 15.3 pA and relatively slow inactivation kinetics (Fig. 2D, E). In cells transfected with siRNA against Piezo1, the current was nearly abolished (an averaged maximal current of 4.4 ± 0.7 pA) (Fig. 2D, E), demonstrating that the recorded mechanically activated current was mediated by Piezo1. Importantly, siRNA-mediated knockdown of either E-cadherin or β-catenin significantly reduced the current, resulting in an averaged maximal current of 45.8 ± 9.6 pA and 35.7 ± 8.7 pA, respectively (Fig. 2D, E).

To examine whether the Piezo1-mediated responses in MDCK cells depend on actin cytoskeleton, we pre-treated the cells with cytochalasin D to disrupt actin fibers. Importantly, we observed significant reduction of both Yoda1-evoked Ca^2+^ response (Fig. 2F, G) and mechanically activated currents (88.7 ± 5.7 pA vs 35.7 ± 6.2 pA for non-treated vs treated cells) (Fig. 2H) in cytochalasin D-treated MDCK cells.

Together, these data show that disruption of either E-cadherin, β-catenin or actin fibers in the mechanotransduction complex all led to impaired Piezo1-mediated functional responses in MDCK cells. Thus, in line with their biochemical interaction, the function of Piezo1 in MDCK cells is profoundly influenced by the E-cadherin-β-catenin-actin mechanotransduction complex, suggesting a robust regulatory mechanism.

### Heterologous interaction of E-cadherin with Piezo1 and Piezo2

To mechanistically understand how Piezo channels are regulated by E-cadherin interaction, we examined their biochemical interaction and functional regulation in heterologous HEK293T cells. Given that both Piezo channels and E-cadherin are localized on the plasma membrane (PM), we first reasoned that their direct interaction might mediate the formation of the macromolecular complex of Piezo1, E-cadherin, β-catenin, vinculin and actin detected in MDCK cells. In line with this, in cells co-transfected with constructs respectively encoding the Piezo1-GST and GFP-E-cadherin fusion proteins, affinity pull-down of the Piezo1-GST proteins using the glutathione-conjugated beads led to co-pull-down of the GFP-E-cadherin proteins, which were detected with western blotting detection using the anti-GFP antibody (Fig. 3A). In contrast, no GFP-E-cadherin proteins were detected from the pull-down sample derived from cells co-transfected with GST and GFP-E-cadherin (Fig. 3A). Similarly, we detected interaction between Piezo2-GST and GFP-E-cadherin as well (Fig. 3B). Thus, in line with the observed endogenous interaction between Piezo1 and E-cadherin in MDCK cells (Fig. 1B, C), heterologously expressed Piezo channels can interact with E-cadherin.

**Figure 3.**
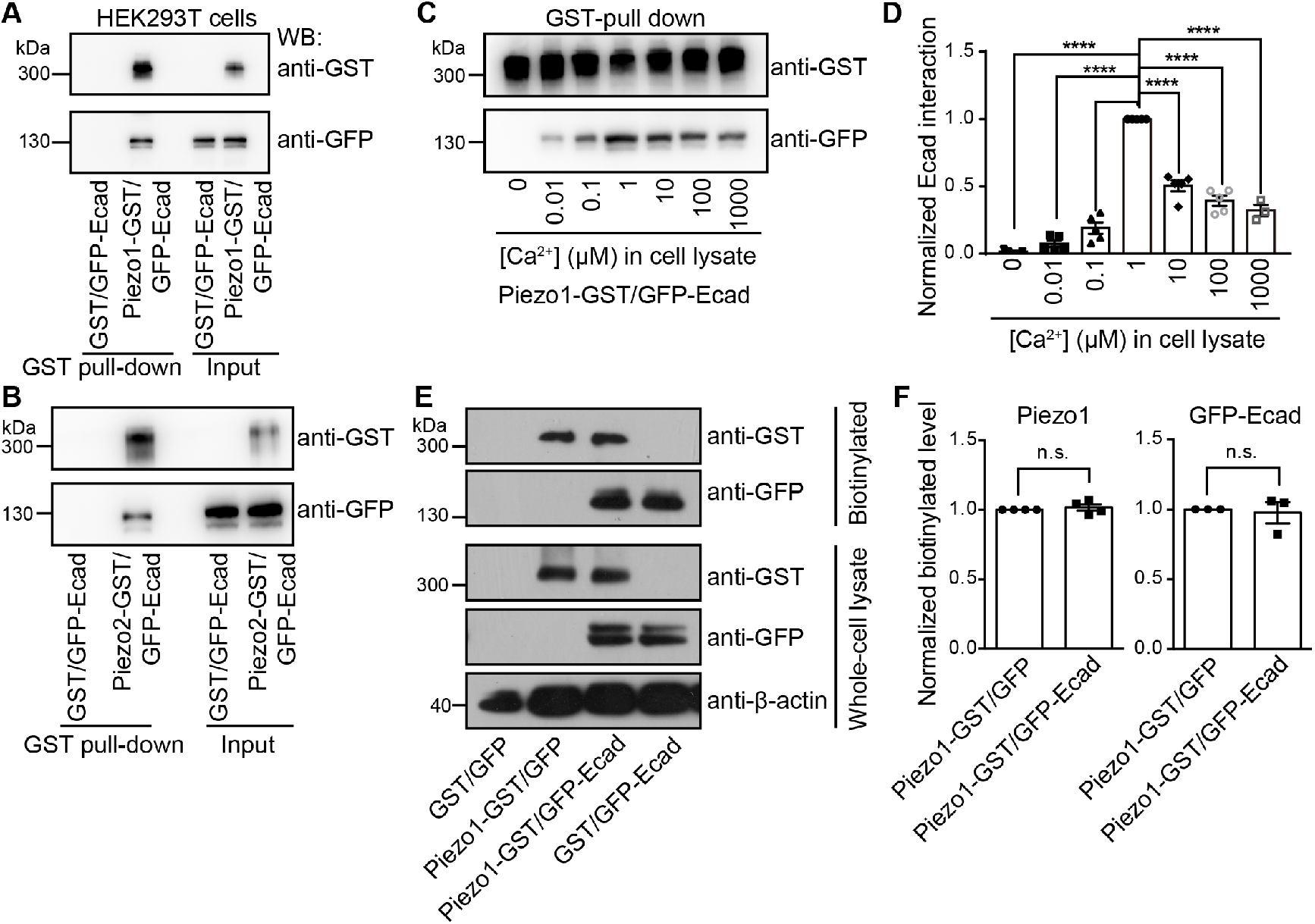
Heterologous interaction of E-cadherin with Piezo1 and Piezo2. (A) Representative Western blots showing GST pull-down of GFP-E-cadherin proteins from HEK293T co-transfected with Piezo1-GST and GFP-E-cadherin, but not from cells co-transfected with GST and GFP-E-cadherin. (B) Representative Western blots showing GST pull-down of GFP-E-cadherin proteins from HEK293T co-transfected with Piezo2-GST and GFP-E-cadherin, but not from cells co-transfected with GST and GFP-E-cadherin. (C) Representative Western blots showing GST pull-down of GFP-E-cadherin proteins from HEK293T co-transfected with Piezo1-GST and GFP-E-cadherin with the indicated Ca^2+^ concentration in the cell lysate aliquots. (D) Scatter plot of normalized interaction between Piezo1-GST and GFP-E-cadherin with the indicated Ca^2+^ concentration. Data are shown as mean ± s.e.m. One-way ANOVA with comparison to the condition of 1 μM Ca^2+^. ****P <0.0001. (E) Representative Western blots of the biotinylated or whole-cell lysate samples derived from HEK293T cells transfected with the indicated constructs. (F) Scatter plots of the normalized level of biotinylated Piezo1 or E-cadherin derived from cells transfected with the indicated constructs. The level of β-actin in the cell lysates was detected using the anti-β-actin antibody for loading control. Unpaired Student’s t-test. n.s. represents non-significance.

Homotypic interactions of E-cadherin with neighboring cells require the presence of millimolar level of extracellular Ca^2+^ (Brasch et al., 2012; Lewis et al., 1994). In line with this, in vitro Ca^2+^ binding assay revealed an average Kd (dissociation constant) of 460 μM to the extracellular EC repeats 1-2 of E-cadherin (Koch et al., 1997). Furthermore, activation of Piezo channels can increase the intracellular Ca^2+^ from a resting level of ~100 nM to ~1 μM. We therefore examined whether the interaction between Piezo1 and E-cadherin depends on Ca^2+^. By varying the Ca^2+^ concentration in cell lysates derived from HEK293T cells co-transfected with Piezo1-GST and GFP-E-cadherin, we observed that the Piezo1-E-cadherin interaction displayed an intriguing bell-shaped dose-dependence of Ca^2+^ (Fig. 3C, D). The interaction was not detected in the absence of Ca^2+^, started to occur in the presence of as low as 10 nM Ca^2+^ and the resting intracellular Ca^2+^ level of 100 nM, peaked at 1 μM Ca^2+^, and then declined with further increasing the Ca^2+^ concentrations to 10 μM, 100 μM and 1 mM (Fig. 3C, D). These data suggest that the Piezo1-E-cadherin interaction does no depend on millimolar level of extracellular Ca^2+^, which is required for homotypic interactions of E-cadherin. In line with this, we observed co-localization of Piezo1 and E-cadherin at cell periphery independently of E-cadherin homotypic interactions with neighboring keratinocytes cultured in medium with 70 μM extracellular Ca^2+^ (Fig. 1E). Instead, the Piezo1-E-cadherin interaction might dynamically depend on the change of intracellular Ca^2+^ varying from 100 nM to 1 μM.

Given that their functional roles depend on their PM expression, we next asked whether the interaction between Piezo1 and E-cadherin might affect their PM expression by carrying out cell surface protein biotinylation assay. Proteins residing in PM were biotinylated and then pulled down via streptavidin-conjugated beads. Based on western blotting of either Piezo1-GST or GFP-E-cadherin in the whole-cell lysates derived from HEK293T cells co-transfected with the recombination of Piezo1-GST/GFP, Piezo1-GST/GFP-E-cadherin, GST/GFP-E-cadherin, we found that co-transfection of Piezo1-GST/GFP-E-cadherin did not affect the overall expression of either Piezo1-GST or GFP-E-cadherin (Fig. 3E). Furthermore, western blotting of the biotinylated samples revealed that the abundance of biotinylated Piezo1-GST or GFP-E-cadherin was not affected by co-expression of these two proteins (Fig. 3E, F), prompting us to hypothesize that E-cadherin might directly affect the mechanosensitivity of Piezo1 for regulating its channel activities.

### Heterologous regulation of Piezo channel function by E-cadherin

We found that the poking-evoked whole-cell current from sparsely re-plated individual HEK293T cells co-expressing Piezo1 and E-cadherin is about 3-fold of that from cells expressing Piezo1 without E-cadherin (6.2 ± 1.1 nA vs 1.9 ± 0.4 nA, respectively) (Fig. 4A, B), indicating a robust potentiation effect of E-cadherin on Piezo1 channel function. Furthermore, the inactivation kinetics of the Piezo1-mediated current was significantly slowed upon co-expression of E-cadherin (Fig. 4A, C). E-cadherin also potentiated the Piezo2-mediated poking-evoked currents (Fig. 4D-F). Given that co-expression of E-cadherin did not affect the abundance of PM-residing Piezo1 (Fig. 3E, F), these data suggest that the enhanced currents recorded from cells co-expressing Piezo1 and E-cadherin is not due to increased PM-expression of Piezo1. To test whether E-cadherin might instead increase the mechanosensitivity of Piezo1, we measured stretch-induced currents by applying negative pressure to the membrane patch under a cell-attached recording configuration. Co-expression of E-cadherin led to a left-shifted pressure-current curve (Fig. 4G), significantly enlarged maximal stretch current (Fig. 4H), and reduced P_50_ value that indicates the pressure required for half maximal activation of Piezo1 (Fig. 4I). These data suggest that E-cadherin potentiates the mechanosensitivity of Piezo1. Additionally, we observed increased Yoda1-induced Ca^2+^ responses in confluent cells co-expressing Piezo1 and E-cadherin (Fig. 4J).

**Figure 4.**
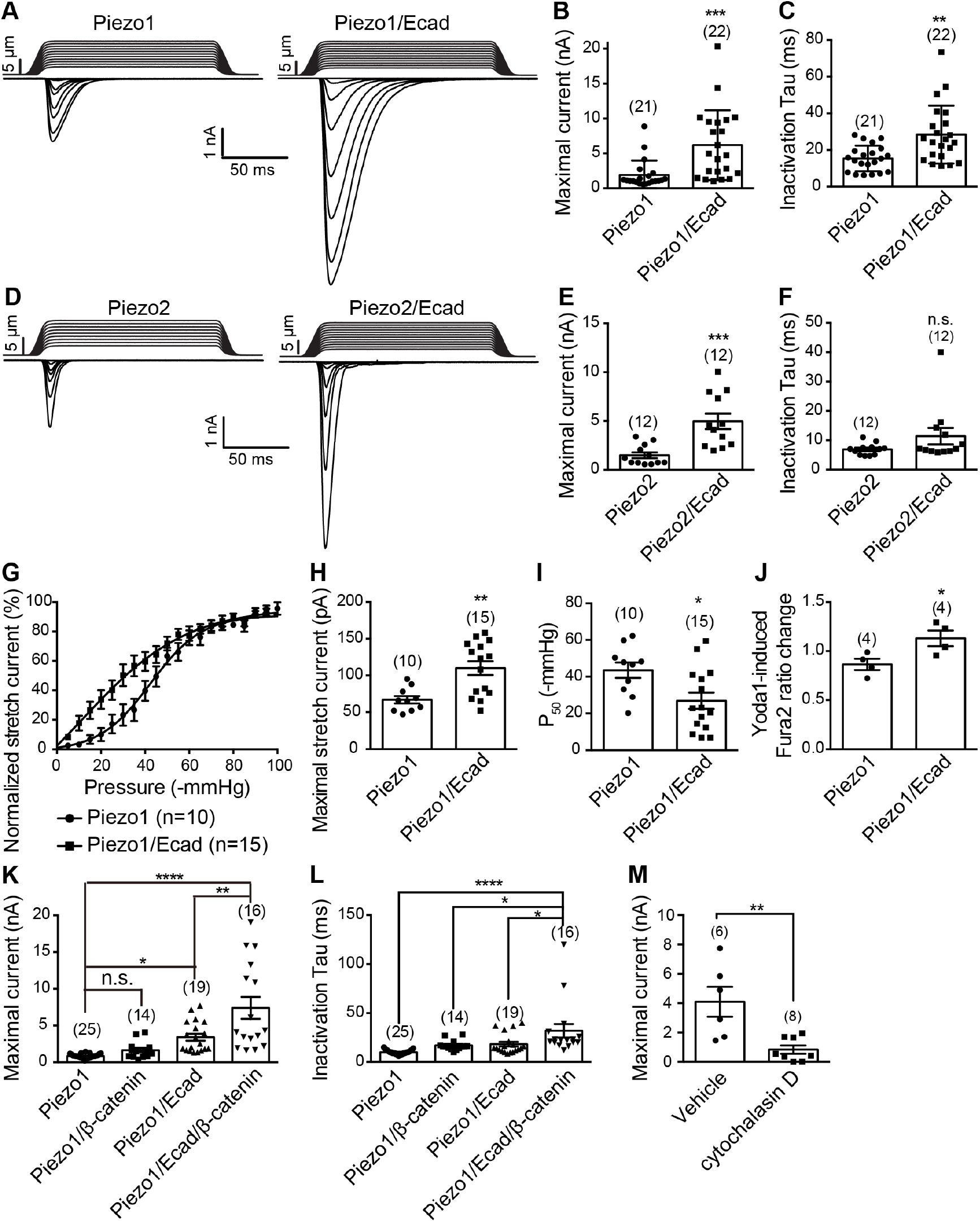
Heterologous regulation of Piezo channel function by E-cadherin. (A, D) Representative poking-evoked whole-cell currents from sparsely re-plated individual HEK293T cells transfected with the indicated constructs. (B, E) Scatter plot of the maximal poking-evoked whole-cell currents from HEK293T cells transfected with the indicated constructs. Each bar represents mean ± s.e.m., and the recorded cell number is labeled above the bar. Unpaired Student’s t-test. ***P < 0.001. (C, F) Scatter plot of the inactivation Tau of poking-evoked whole-cell currents from HEK293T cells transfected with the indicated constructs. Each bar represents mean ± s.e.m., and the recorded cell number is labeled above the bar. Unpaired Student’s t-test. **P < 0.01. (G) Pressure-current response curves from HEK293T transfected with the indicated constructs. The curves were fitted with a Boltzmann equation. (H) Scatter plot of the maximal stretch-induced current from HEK293T cells transfected with the indicated constructs. Each bar represents mean ± s.e.m., and the recorded cell number is labeled above the bar. Unpaired Student’s t-test. **P < 0.01. (I) Scatter plot of the P_50_ values calculated from individual cells transfected with the indicated constructs. Each bar represents mean ± s.e.m., and the recorded cell number is labeled above the bar. Unpaired Student’s t-test. *P < 0.05. (J) Scatter plot of Yoda1-induced Fura-2 ratio change of confluent HEK293T cells transfected with the indicated constructs. Each bar represents mean ± s.e.m., and the imaged coverslip number is labeled above the bar. Unpaired Student’s t-test. *P < 0.05. (K, L) Scatter plot of the maximal poking-evoked whole-cell currents (K) and inactivation Tau (L) from HEK293T cells transfected with the indicated constructs. Each bar represents mean ± s.e.m., and the recorded cell number is labeled above the bar. One-way ANOVA with multiple comparison test. ****P<0.0001, **P < 0.01, *P < 0.05. (M) Scatter plot of the maximal poking-evoked whole-cell currents from HEK293T cells transfected with the indicated constructs pre-treated with either vehicle or 10 μM cytochalasin D. Each bar represents mean ± s.e.m., and the recorded cell number is labeled above the bar. Unpaired Student’s t-test. **P < 0.01.

When compared to currents recorded from cells co-expressing Piezo1 and E-cadherin, overexpression of β-catenin together with Piezo1 and E-cadherin led to an additional increase of poking-evoked current (Fig. 4K) and further slowing of the inactivation kinetics (Fig. 4L). However, co-expression of Piezo1 and β-catenin did not cause potentiation of the Piezo1 current (Fig. 4K, L), indicating the requirement of E-cadherin for regulation. Furthermore, we found that cytochalasin D-treatment of HEK293T cells co-expressing Piezo1 and E-cadherin led to about 80% reduction of the poking-evoked current (4.1 ± 1.0 nA vs 0.8 ± 0.3 nA for vehicle- or cytochalasin D-treated cells, respectively) (Fig. 4M), supporting the involvement of cytoskeleton in E-cadherin-mediated regulation of Piezo1. Thus, in line with the observation in MDCK cells, these data demonstrate that Piezo1 is regulated through the E-cadherin-β-catenin-actin mechanotransduction complex.

### Identifying the interacting domains between Piezo1 and E-cadherin

We next went on to identify the interaction region between Piezo1 and E-cadherin. E-cadherin is a single-pass transmembrane protein containing a N-terminal ectodomain (ED) composed of five tandem EC repeats, which are involved in homotypic interaction between neighboring cells, and a C-terminal cytoplasmic tail (CT) for interaction with β-catenin and other cytoplasmic proteins (Fig. 5A). To identify which region of E-cadherin might interact with Piezo1, we generated Flag-tagged full-length E-cadherin or its fragments respectively containing the ED, the CT, and the transmembrane domain (TM) together with the CT (TM-CT). We found that Piezo1-GST was able to pull down the co-expressed full-length E-cadherin, the ED and the TM-CT, but not the CT alone (Fig. 5B), suggesting that multiple regions of E-cadherin participate in interaction with Piezo1.

**Figure 5.**
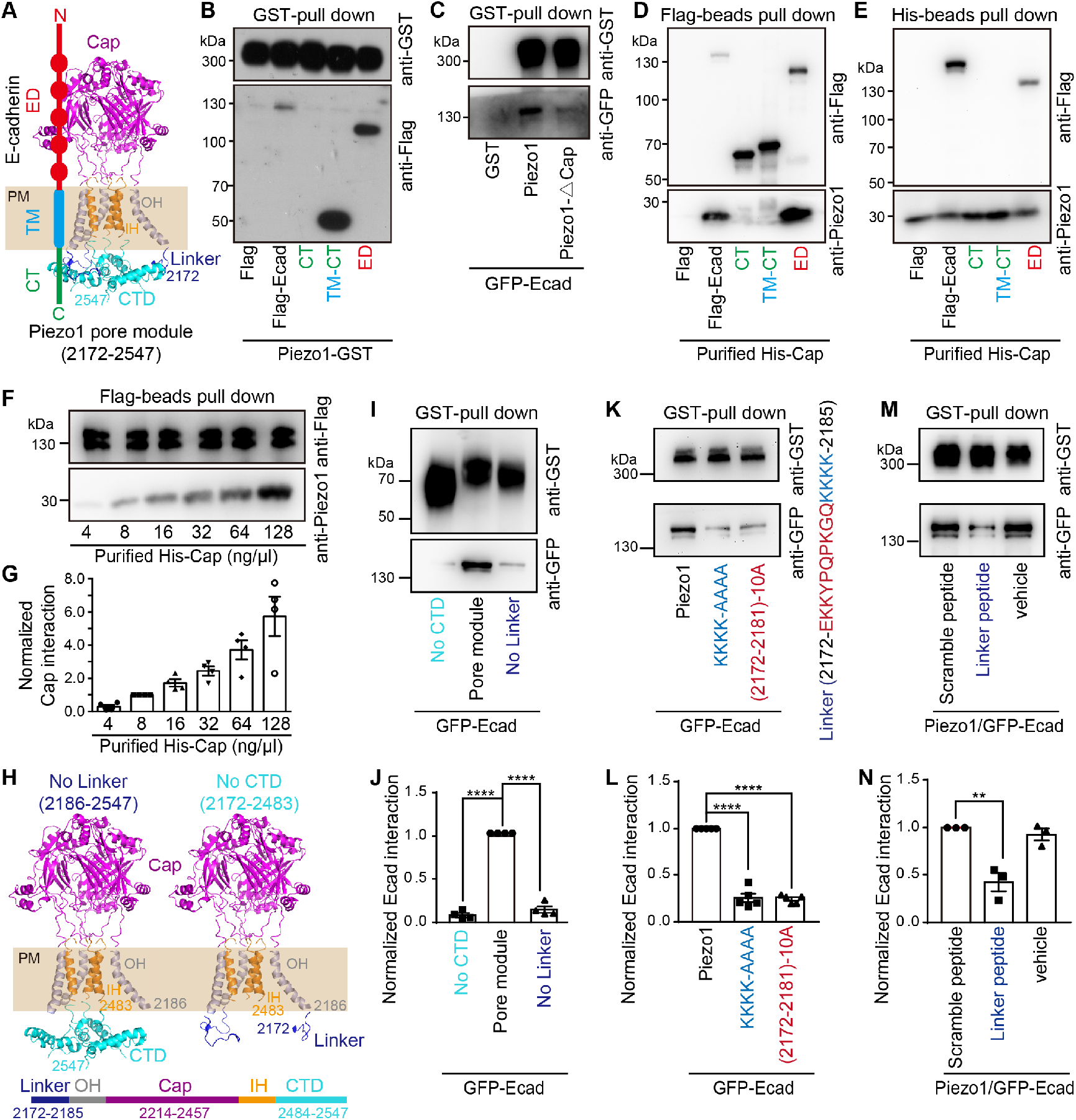
Identifying the interacting domains between Piezo1 and E-cadherin. (A) Diagram showing the central pore module of Piezo1 containing the linker (in blue), outer helix (OH, gray), the extracellular cap domain (in purple), the pore-lining inner helix (IH, in orange), the intracellular C-terminal domain (CTD, in cyan), and the topological representation of E-cadherin containing the N-terminal ectodomain (ED), the single transmembrane region (TM) and the intracellular C-terminal tail (CT). (B) Representative Western blots showing GST pull-down of the Flag-tagged full-length E-cadherin or the indicated fragment proteins from HEK293T cells co-transfected with Piezo1-GST and the indicated E-cadherin constructs. (C) Representative Western blots showing GST pull-down of GFP-E-cadherin from HEK293T cells co-transfected GFP-E-cadherin with either GST, Piezo1-GST or Piezo1-ΔCap. (D) Representative Western blots showing that anti-Flag antibody immunoprecipitation of the purified Cap fragment protein, which was incubated with cell lysates derived from HEK293T cells transfected with the indicated full-length or fragment E-cadherin. The Cap fragment protein was detected using the Piezo1 antibody that was raised against the Cap fragment protein. (E) Representative Western blots showing that His-beads pull-down of the Flag-tagged full-length E-cadherin or the indicated fragment proteins using the purified His-tagged Cap fragment proteins. (F) Representative Western blots showing that dose-dependent pull-down of the purified His-tagged Cap fragment protein by the Flag-E-cadherin protein. (G) Scatter plot of normalized interaction of purified His-tagged Cap fragment protein with the Flag-E-cadherin protein. Data are shown as mean ± s.e.m. (H) Diagram showing the pore module lacking either the linker region (residues 2186-2547) or the CTD (residues 2172-2483). (I, K) Representative Western blots showing GST pull-down of GFP-E-cadherin proteins from HEK293T cells co-transfected with GFP-E-cadherin and the indicated Piezo1 mutant constructs. (M) Representative Western blots showing GST pull-down of GFP-E-cadherin proteins from HEK293T cells co-transfected with GFP-E-cadherin and Piezo1-GST with the indicated treatment conditions. (J - N) Scatter plot of the normalized interacting E-cadherin level shown in I, K, M, respectively. Data shown as mean ± s.e.m. One-way ANOVA with multiple comparison test. ****P <0.0001, **P < 0.01.

The observation that the extracellular ED of E-cadherin is involved in interacting with Piezo1 led us to reason whether the large extracellular cap domain of Piezo1 might serve as the reciprocal binding region (Fig. 1A). In line with this idea, deleting the cap domain resulted in the mutant protein Piezo1-ΔCap with reduced ability to pull down E-cadherin (Fig. 5C), despite that Piezo1-ΔCap proteins remain the ability to form trimeric architectures (Ge et al., 2015). We further purified the His-tagged Cap fragment protein (Ge et al., 2015) and examined its ability to interact with either the full-length E-cadherin or the E-cadherin fragments. The purified Cap fragment protein specifically interacted with the full-length E-cadherin and the ED, but not the CT or the TM-CT (Fig. 5D, E). Furthermore, the full-length E-cadherin was able to pull down the purified Cap fragment protein in a dose-dependent manner (Fig. 5F, G). These data demonstrate that the extracellular ED of E-cadherin directly interacts with the extracellular cap domain of Piezo1.

The cap domain constitutes the extracellular structure of the central pore module (residues 2172-2547), which also contains the intracellular C-terminal domain (CTD) (residues 2484-2547) and is preceded with a 14-residue-constituted linker region (residues 2172-2185) that connects the anchor domain to the outer helix of the pore module (Fig. 5A, H and Fig. 7B). We have previously found that the linker, but not the CTD, is required for interaction with the sarco-endoplasmic reticulum ATPase (SERCA) (Zhang et al., 2017). We thus examined the interaction of E-cadherin with the pore module fragment containing residues 2172 to 2547 (Fig. 5A), the pore module lacking the linker (residues 2186-2547) (Fig. 5H) or the pore module without the CTD (residues 2172-2483) (Fig. 5H). The pore module was able to sufficiently interact with E-cadherin (Fig. 5I). Interestingly, deleting either the linker or the CTD drastically reduced the ability of the pore module fragment to pull down the co-expressed GFP-E-cadherin (Fig. 5I, J), suggesting that both the linker and the CTD are required for E-cadherin binding to the pore module. We further examined the involvement of the 14-residue-constituted linker in mediating the interaction between the full-length Piezo1 and E-cadherin using the previously generated Piezo1-(2172-2181)10A-GST (residues 2172-2181 were mutated to alanine) and Piezo1-KKKK-AAAA-GST [the cluster of 4 lysine residues (2182-2185) was mutated to alanine] mutants (Zhang et al., 2017) (Fig. 5K). Both mutants had significantly reduced ability to pull down the co-expressed E-cadherin (Fig. 5K, L). Given that the linker region is critically required for E-cadherin interaction to both the full-length Piezo1 and the C-terminal pore module fragment, we reasoned that it might serve as a direct binding site. In line with this, we found that application of the synthesized membrane-permeable linker-peptide (Zhang et al., 2017) to cell lysates derived from HEK293T cells co-expressing Piezo1 and E-cadherin significantly reduced Piezo1-E-cadherin interaction, while the scramble peptide- or vehicle-treated samples had no such effect (Fig. 5M, N), indicating that the linker-peptide and Piezo1 competed for E-cadherin interaction. Collectively, these data suggest that the linker region might serve as a direct binding site for the intracellular region of E-cadherin. The close proximity of the linker near the PM (Fig. 5A, H) indicates that the reciprocal intracellular binding region in E-cadherin might need to be tethered to the plasma membrane for interaction, which might explain the PM-targeting TM-CT fragment but not the cytosolic CT fragment of E-cadherin, robustly bind with Piezo1 (Fig. 5B). Taken together, these data demonstrate that multiple regions in both Piezo1 and E-cadherin are involved in their interaction.

### Functional regulation of Piezo1 by E-cadherin through the cap and the linker

We next examined whether E-cadherin might functionally regulate Piezo1 through the identified interacting domains in Piezo1 including the intracellular linker and the extracellular cap, both of which are critical for mechanogating of Piezo1 (Wang et al., 2019; Zhang et al., 2017). While co-expression of E-cadherin consistently enhanced Piezo1-mediated currents, it did not cause significant potentiation of the poking-evoked currents mediated by the linker mutants Piezo1-(2172-2181)10A and Piezo1-KKKK-AAAA (Fig. 6A, B). In line with our previous report that the linker is critical for mechanogating (Zhang et al., 2017), Piezo1-(2172-2181)10A and Piezo1-KKKK-AAAA had reduced mechanically activated currents (Fig. 6A, B). Furthermore, application of the membrane-permeable linker peptide to HEK293T cells co-expressing Piezo1 and E-cadherin significantly prevented the potentiated currents (Fig. 6C). As a control, the scramble peptide had no such effect (Fig. 6C). Together with the finding that the linker peptide disrupted the biochemical interaction between Piezo1 and E-cadherin (Fig. 5M, N), these data provide strong evidence that the linker is directly involved in E-cadherin regulation of Piezo1. Intriguingly, previous studies have shown that the linker is involved in SERCA2 binding and inhibition of Piezo1 (Zhang et al., 2017). The opposite role of E-cadherin and SERCA2 in affecting Piezo1 function could be explained by the differential involvement of the CTD in binding of E-cadherin and SERCA2 to Piezo1. While CTD is required for E-cadherin binding to Piezo1 (Fig. 5I, J), it prevents the binding of SERCA2 (Zhang et al., 2017). These analyses suggest that the linker and CTD might work together to mediate the potentiation effect of E-cadherin on Piezo1 via the cytosolic interaction.

**Figure 6.**
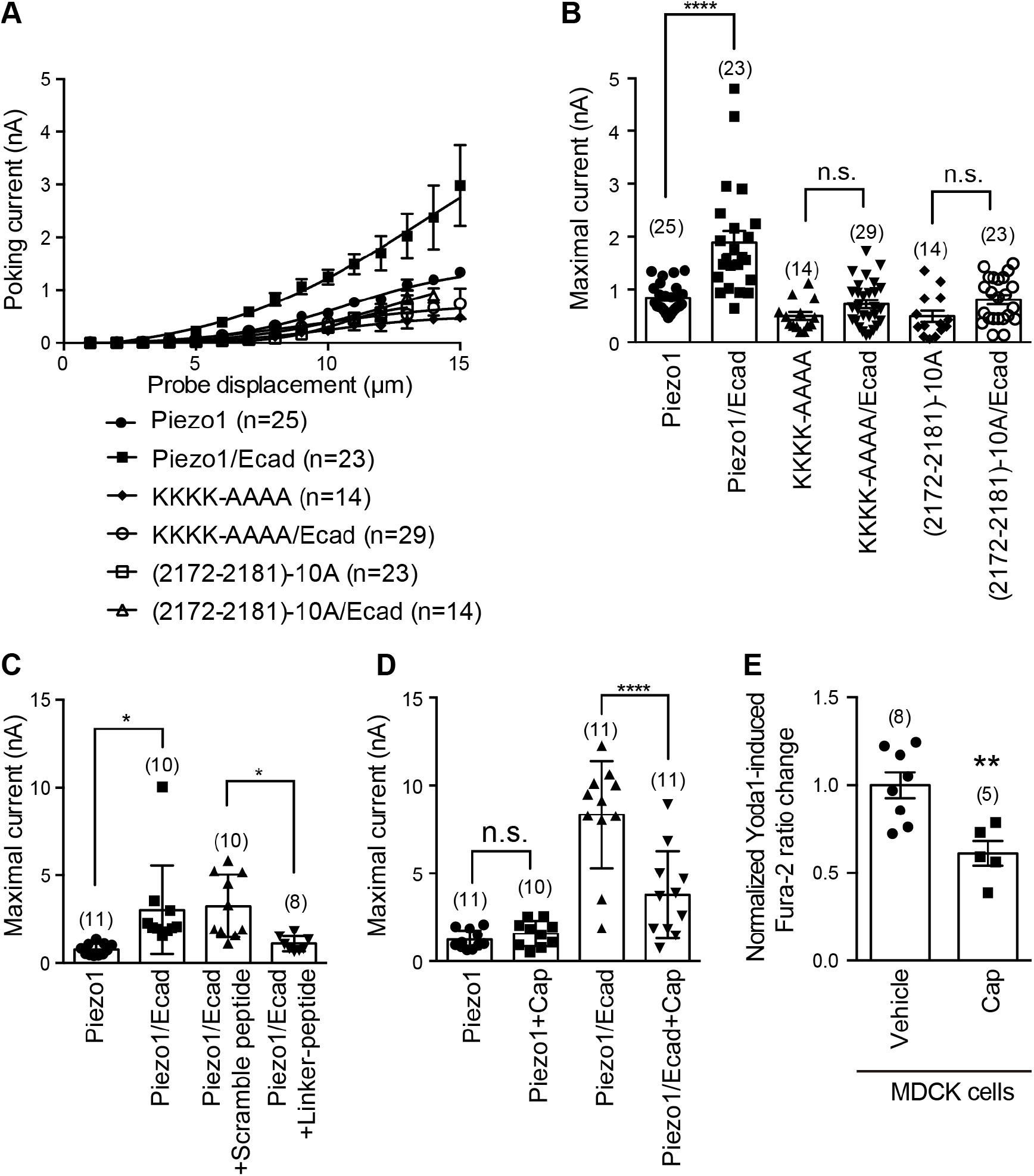
Regulation of Piezo1 by E-cadherin through the cap and the linker domains. (A) Probe displacement and current relationship of HEK293T cells transfected with the indicated constructs. The number of recorded cells are labeled. (B) Scatter plot of the maximal poking-evoked whole-cell currents from HEK293T cells transfected with the indicated constructs. Each bar represents mean ± s.e.m., and the recorded cell number is labeled above the bar. Unpaired Student’s t-test. ***P < 0.001. (C) Scatter plot of the maximal poking-evoked whole-cell currents from HEK293T cells transfected with the indicated constructs and treated with either the membrane-permeable scramble or the Linker-peptide. Each bar represents mean ± s.e.m., and the recorded cell number is labeled above the bar. Unpaired Student’s t-test. *P < 0.05. (D) Scatter plot of the maximal poking-evoked whole-cell currents from HEK293T cells transfected with the indicated constructs and treated with or without the purified Cap fragment proteins. Each bar represents mean ± s.e.m., and the recorded cell number is labeled above the bar. Unpaired Student’s t-test. ****P < 0.0001. (E) Scatter plot of normalized Yoda1-induced Fura-2 ration change from MDCK cells treated with vehicle or the purified Cap fragment protein. Each bar represents mean ± s.e.m., and the number of imaged coverslips is labeled above the bar. Unpaired Student’s t-test. P < 0.01.

Given the critical role of the cap in mediating interaction with E-cadherin, we reasoned that the purified Cap fragment protein might disrupt regulation of Piezo1 by E-cadherin. Indeed, application of the purified Cap protein into the extracellular recording buffer significantly reduced the poking-evoked currents in HEK293T cells co-expressing Piezo1 and E-cadherin (Fig. 6D). In contrast, such treatment did not cause any current reduction in cells expressing Piezo1 without E-cadherin (Fig. 6D). These data demonstrate a specific regulatory effect of the Cap-E-cadherin binding on Piezo1 channel function. To examine whether the Cap domain might also participate in endogenous regulation of Piezo1 by E-cadherin, we applied the purified Cap protein to MDCK cells and found a significant reduction of Yoda1-induced Ca^2+^ response in the presence of the Cap protein (Fig. 6E). Together, these data demonstrate that both heterologous and endogenous regulation of Piezo1 channel function by E-cadherin requires the cap domain.

## Discussion

The mechanically activated Piezo channel, including Piezo1 and Piezo2 (Coste et al., 2010; Coste et al., 2012), represent a class of versatile mechanotransducers for mediating distinct forms of mechanical stimuli in a wide variety of cell types and consequently governing a broad spectrum of biological processes (Douguet and Honore, 2019; Murthy et al., 2017; Wu et al., 2016; Xiao, 2019). Elucidating its mechanogating mechanism is fundamental to understand how it might fulfill its mechanotransduction function in intact cells. Mechanosensitive ion channels are generally gated by mechanical forces via either the force-from-lipids model or the force-from-filament model (or tether model) (Anishkin et al., 2014). Here we show biochemical and functional findings to demonstrate that Piezo1 is physically linked to the actin cytoskeleton via the E-cadherin-β-catenin-vinculin mechanotransduction complex and the direct interaction of E-cadherin to key mechanogating domains of Piezo1, for the first time establishing a force-from-filament or tether model for the Piezo channel. In line with this gating mechanism, previous studies have shown that pharmacological disruption of the actin cytoskeleton impairs Piezo1-mediated currents evoked by cell indention (Gottlieb et al., 2012) or elastomeric pillar deflection at cell-substrate contact points (Bavi et al., 2019).

Several lines of evidence suggest that the regulatory effect of E-cadherin on Piezo1 might not depend on E-cadherin’s homotypic interactions between neighboring cells. First, the co-localization of Piezo1, E-cadherin and F-actin in keratinocytes was detected in plasma membrane locations free of cell-cell contacts (Fig. 1E). Second, Piezo1-E-cadherin interactions do not depend on high concentration of Ca^2+^, which is required for E-cadherin’s homotypic interactions (Fig. 3C, D). Third, functional potentiation of Piezo1-mediated currents by E-cadherin could be detected in individual cells without contacting cells and in cell-attached membranes (Fig. 2D, E and 4A-I). Together, these data suggest that E-cadherin-mediated adhesions junctions might not be necessarily needed for the tethered gating of Piezo channels. Instead, F-actin-derived force transduction might modulate the mechanosensitivity of Piezo channels. In line with this, endogenous traction forces have been reported to locally activate Piezo1 (Ellefsen et al., 2019; Pathak et al., 2014).

On the basis of the identification of the transmembrane gate located in the inner helix (IH) (Wang et al., 2019) and the cytosolic lateral plug gates that physically block the three lateral ion-conducting portals (Geng et al., 2020), we have proposed that Piezo channels might utilize a dual-gating mechanism, in which the transmembrane gate is dominantly controlled by the top extracellular cap (Wang et al., 2019), while the lateral plug gates are controlled by the peripheral blade-beam apparatus via a plug-and-latch mechanism (Geng et al., 2020; Wang et al., 2019; Wang et al., 2018; Zhao et al., 2018a; Zhao et al., 2018b) (Figure 7A, B). The cap domain is embedded in the center of the nano-bowl shaped by the curved Piezo1-membrane system and sits right on the top of the transmembrane pore of Piezo1 (Fig. 7A, B). Structural analysis has revealed that motion of the cap domain is strictly coupled to the transmembrane gate residing in the pore-lining inner helix (Fig. 7A, B) (Wang et al., 2019). Furthermore, either deleting (Wang et al., 2019) or crosslinking the cap (Lewis and Grandl, 2020) completely abolished mechanical activation of Piezo1. These studies have led us to propose that the cap domain functions as a critical mechanogating domain to predominantly control the transmembrane gate (Wang et al., 2019). Following the transmembrane pore, Piezo channels utilize three lateral portals equipped with threes splicable lateral plug gates as their intracellular ion-conducting routes (Geng et al., 2020) (Fig. 7B). Interestingly, the three lateral plug gates physically block the lateral portals and are strategically latched onto the central axis for coordinated gating (Geng et al., 2020) (Fig. 7B). Remarkably, we have identified that the extracellular ectodomain of E-cadherin directly interacts with the extracellular cap domain of Piezo1, while its cytosolic tail with the TM region might interact with the cytosolic linker and CTD of Piezo1 (Fig. 5, and Fig. 6), which are in close proximity to the lateral plug gates. Thus, a physical interaction of E-cadherin with Piezo1 might allow a direct focus of cytoskeleton-transmitted force on the extracellular cap domain and the intracellular linker and CTD to gate the transmembrane gate and cytosolic lateral plug gates, respectively (Fig. 7A, B), providing a structural basis for a tether model for mechanogating of both gates.

**Figure 7.**
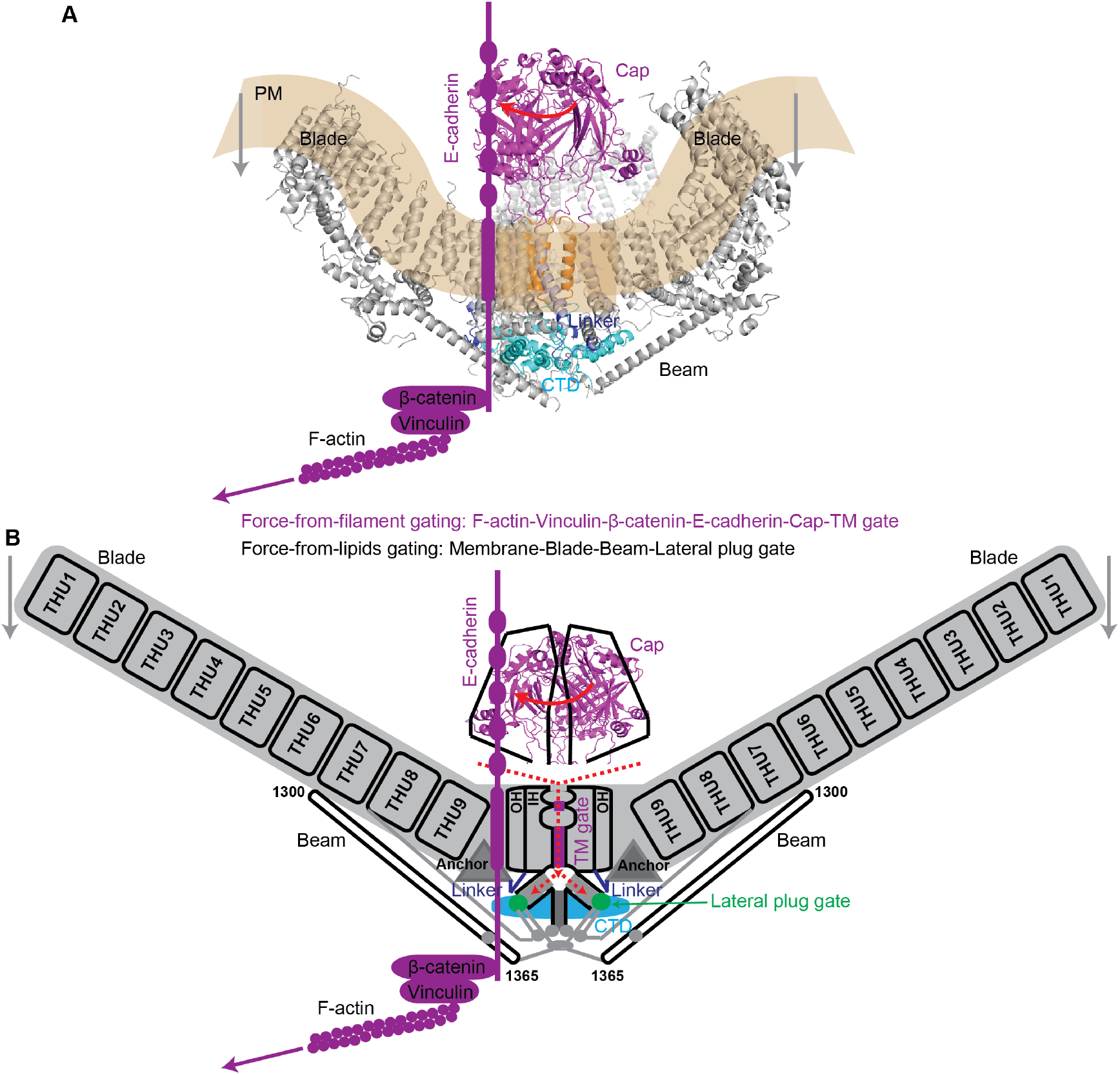
A proposed mechanogating model for Piezo channel integrating both the E-cadherin-mediated tether model and the intrinsic force-from-lipids model. The Piezo1 structure (A) or its depicted model showing two out of its three subunits (B) in complex with E-cadherin, β-catenin, vinculin and F-actin. The featured structural domains including the blade with 36 TMs, which are folded into 9 repetitive transmembrane helical units (THU) consisting 4 TMs each, the long intracellular beam, the extracellular cap (in purple), the linker (in blue) that connecting the anchor domain to the outer helix (OH), and the intracellular C-terminal domain (CTD, in cyan) following the inner helix (IH) are labeled in both (A) and (B). The transparent yellow area in (A) indicates the nano-bowl-shaped plasma membrane curved by the highly non-planar transmembrane blade of the Piezo1 structure. The Piezo1 model in (B) additionally depicts the blade composed of the THU1 to THU9, the transmembrane gate (TM gate, in purple) represented by the constriction sites along the transmembrane pore enclosed by the IHs, the lateral ion-conducting portals following the transmembrane pore, and the lateral plug gates shown in green spheres that physically block the outlets of the lateral portals. The ion-permeating pathway is indicated by the dashed red arrow. The interactions between the ectodomain of E-cadherin with the cap domain of Piezo1, and the cytosolic tail of E-cadherin with the linker and CTD of Piezo1 constitute the structural basis for a tether model of gating the TM gate and cytosolic lateral plug gates illustrated in (B). Membrane tension-induced flattening of the highly curved transmembrane blades and the residing membrane might constitute the structural basis for a force-from-lipids model of gating the cytosolic lateral plug gates via the blade-beam-lateral plug gate transduction pathway. The two gating models might synergistically gate the lateral plug gates via converge at the intracellular linker and CTD, which are in close contact with the lateral plug gate. The red arrow indicates the rotation of the cap, which is coupled to the TM gate. The grey arrow indicates membrane tension, which can lead to flattening of the blade. The purple arrow indicates force transmitted by the actin-cytoskeleton.

In addition to the tether model revealed in the present study, the Piezo channel might utilize its signature bowl-shaped structural feature and intrinsic mechanosensitivity to adopt the force-from-lipids model for mechanogating. Structural analysis reveals large conformational changes of the highly curved blades, which might be converted to the cytosolic lateral plug gates via a lever-like motion of the featured beam structure (Fig. 7A, B) (Geng et al., 2020; Wang et al., 2019; Wang et al., 2018; Zhao et al., 2018a). Using the probe of an atomic force microscope to apply force to the Piezo1 channel reconstituted in lipid bilayers, Lin et al. have elegantly demonstrated that the highly curved blade can undergo reversible flattening at biologically relevant pressures (Lin et al., 2019). Furthermore, we have identified a serial of key mechanotransduction sites along the blade-beam-lateral plug gate (Geng et al., 2020; Wang et al., 2018; Zhao et al., 2018a), which might constitute an intramolecular mechanotransduction pathway for converting long-range conformational changes from the distal blades to the cytosolic lateral plug gates (Fig. 7B) (Geng et al., 2020; Wang et al., 2018; Xiao, 2019; Zhao et al., 2018a). Collectively, these studies suggest that the blade-beam apparatus might serve as the molecular basis for a force-from-lipids model to gate the cytosolic lateral plug gates (Fig. 7A, B).

Given the existence of the structurally and functionally distinct molecular bases for the force-from-filament and force-from-lipids models, we propose that the Piezo channel might incorporate these two distinct yet non-exclusive gating models to serve as a versatile mechanotransducer (Figure 7A, B). Using the dual mechanogating models, Piezo channels are not only able to constantly monitor changes in local membrane curvature and tension, but also overcome the obstacle of limited membrane tension propagation within an intact cellular membrane to effectively detect long-range mechanical perturbation across a cell or between cells. Indeed, Piezo1 can mediate both localized and whole-cell mechanical responses regardless whether the mechanical stimuli are either exogenously applied or endogenously originated (Bavi et al., 2019; Coste et al., 2010; Cox et al., 2016; Ellefsen et al., 2019; Lewis and Grandl, 2015; Shi et al., 2018).

The dual mechanogating model might also allow Piezo channels to flexibly tune to variable cellular context and to distinct forms of mechanical stimuli. For instance, the two gating models might act synergistically to sensitize the mechanosensitivity of Piezo channels. In line with this, the mechanosensitivity of Piezo1 was measured higher in intact cell membranes (T_50_ of ~1.4 mN/m) (Lewis and Grandl, 2015) than in artificial lipid bilayers (T_50_ of ~3.4 mN/m) (Syeda et al., 2016) and membrane blebs that are presumed to lack actin cytoskeleton (T_50_ of ~4.5 mN/m) (Cox et al., 2016). Importantly, the convergence of the tether model and force-from-lipids model at the cytosolic lateral plug gates might allow a synergistic gating. The two models might also be preferentially utilized in a cellular context-dependent manner. For instance, it has been reported that actin-directed force rather than membrane tension might act dominantly to activate Piezo1 at the cell-substrate interface (Bavi et al., 2019). In contrast, localized perturbation of membrane tension might mainly activate Piezo1 via the force-from-lipids model (Cox et al., 2016; Shi et al., 2018).

The identification of a direct interaction between the Piezo channel-based and the E-cadherin-based mechanotransduction systems might open a novel avenue in our understanding of cellular mechanotransduction in general. While here we have focused on E-cadherin to illustrate the proof-of-principal for the tethered gating model of the Piezo channel, cadherins constitute a large superfamily of adhesion receptors, which are widely expressed in epithelial, endothelial, and nervous system (Niessen et al., 2011). Given that the extracellular cadherin domain of E-cadherin is a shared common structural motif among different cadherin members and is involved in interaction with Piezo1, it is reasonable to propose that other types of cadherins might also regulate Piezo channel function, consequently expanding the cadherin-dependent tether model of Piezo channels to other cell types and systems. Furthermore, the demonstration of a physical interaction between Piezo channels and E-cadherin raises the possibility that Piezo channels might also impact E-cadherin-mediated mechanotransduction process via at least two foreseeing mechanisms: Piezo channel-mediated Ca^2+^ signaling [e.g. the observed bell-shaped dose-dependence of Piezo1-E-cadherin interaction on Ca^2+^ (Fig. 3C, D)] and a reciprocal force transmission from Piezo channels to the E-cadherin-mediated mechanotransduction complex. For instance, membrane tension-induced conformational changes of the bowl-shaped Piezo-membrane system might disturb the embedded cap domain and consequently the ectodomain of E-cadherin, which is involved in both trans and cis dimerization of E-cadherin and the formation of adhesions junctions (Leckband and de Rooij, 2014). Future studies will investigate how the two prominent mechanostranduction systems are integrated to regulate cellular mechanotransduction processes.

## Supporting information

The key resource table

## Acknowledgements

We thank Dr. Ardem Patapoutian from the Scripps Research for sharing the Piezo1 and Piezo2 constructs; Dr. Jie Na from Tsinghua University for sharing the canine E-cadherin and β-catenin constructs; Jinyu Wang at the Imaging Core Facility, Technology Center for Protein Sciences at Tsinghua University for assistance with Nikon A1 confocal microscopy. This work was supported by the National Natural Science Foundation of China (31825014 and 31630090) and the National Key R&D Program of China (2016YFA0500402) to B.X.

## Author Contributions

J.W. did biochemical and immunostaining studies and data analysis; J.J. did electrophysiology and Ca^2^+ imaging and data analysis; X.Y. purified the Cap fragment protein; L.W. helped biochemical studies; B.X. conceived and directed the study, and wrote the manuscript with help from all other authors.

## Declare of Interests

The authors declare no competing financial interests.

## STAR Methods

### Key Resources Table

#### Lead Contact and Materials Availability

Further information and requests for resources and reagents should be directed to and will be fulfilled by the Lead Contact, Bailong Xiao. All unique reagents generated in this study are available from the Lead Contact with a completed Materials Transfer Agreement.

### Experimental Model and Subject Details

#### Animals

Naïve male C57BL/6J mice at 8-week-old age were used for Piezo1.1 identification in different tissues. All animal care and procedures were approved and performed in accordance with the Institutional Animal Care and Use Committee (IACUC) guidelines set up by Tsinghua University. All animals were housed in isolated ventilated cages (maximal six mice per cage) barrier facility under conditions of a 12/12-hour light/dark cycle, 22-26 °C with sterile pellet food and water ad libitum in the Tsinghua University animal research facility, which is accredited by the Association for Assessment and Accreditation of Laboratory Animal Care International (AAALAC).

#### Cell lines and Primary Cultured Cell

Human embryonic kidney 293T (HEK293T) cells purchased from ATCC were used for heterologous transfection, biochemistry, electrophysiology, calcium imaging. MDCK cells were generously provided by Dr. Xu Tan at the Tsinghua University. Primary mouse keratinocyte was cultured from newborn mice (P0-P2) according to the previous protocol (Liu et al., 2019). All cells described above were maintained at 37°C with 5% CO_2_.

### Method Details

#### cDNA clones and molecular cloning

The mouse Piezo1 and Piezo2 constructs were generously provided by Dr. Ardem Patapoutian from the Scripps Research, while the canine GFP-E-cadherin and β-catenin constructs were generously provided by Dr. Jie Na from Tsinghua University. The Piezo1-pp-GST-IRES-GFP (Piezo1-GST) and Piezo1-pp-GST-IRES-GFP (Piezo2-GST) constructs were previously generated and used(Ge et al., 2015; Wang et al., 2019; Zhao et al., 2018a). Constructs expressing the E-cadherin fragments, including N-terminal ectodomain of residues 1-711 (ED), the cytosolic C-terminal tail of residues 735-886 (CT), the transmembrane region together with the CT of residues 712-886 (TM-CT) were constructed using the one step cloning kit according to the instruction manual (Vazyme Biotech). All constructs were verified by sequencing. The primers used for generating the constructs are listed in the key resource table.

#### Cell culture and transfection

MDCK cells were cultured in Dulbecco’s Modified Eagle Medium (DMEM) supplemented with 10% fetal bovine serum (FBS), 1% penicillin and streptomycin (P/S, 100 units/ml penicillin and 10 μg/ml streptomycin), and transfected with Lipofectamine 2000 (Invitrogen) according to the manufacturer’s instruction. HEK293T cells were cultured in DMEM supplemented with 10% FBS and 1% P/S, and transfected with polyethylenimine (PEI) (Polysciences).

Primary mouse keratinocytes from newborn mice (P0-P2) was cultured according to the previously described protocol (Liu et al., 2019). Newborn mice were decapitated and soaked into 10% povidone, 70% ethanol and DPBS for 5 min respectively. The back skin was removed, stripped of fat and floated (epidermis upward) in 0.25% trypsin for 1 h at 37°C. Then epidermal layer of the skin was peeled off, cut into smaller pieces in CnT2 medium with 0.07 mM Ca^2+^ (CELLnTEC) supplemented with 10% FBS and 1% P/S, and then stirred for 30 min at room temperature in a 100 ml bottle to get single cells. The cells were filtered with a 70 μm cell strainer, spin down (1000 rpm, 3 min) and resuspended in CnT2 medium without extra serum and P/S. Then cells were plated on coverslips coated with 10 μg/ml fibronectin and 10 μg/ml collagen.

#### Antibodies

The Piezo1 antibody raised against the cap domain (or the C-terminal extracellular domain) of mouse Piezo1 (amino acids 2218-2453) was custom generated by Abgent (Suzhou, China) and previously characterized and used (Zhang et al., 2017). The Piezo1 antibody was used at a concentration of 1:1000 for western blotting. Other antibodies used for western blotting include rabbit anti-GST (Cell Signaling Technology, 1:1000), mouse anti-GFP(Abgent, 1:1000), rabbit anti-vinculin (Cell Signaling Technology, 1:1000), rabbit anti-β-catenin (Cell Signaling Technology, 1:1000), mouse anti-E-cadherin (Abcam, 1:1000), rabbit anti-β-actin (Cell Signaling Technology, 1:1000).

#### GST-pull-down and co-immunoprecipitation

For co-immunoprecipitation of endogenous Piezo1 and adhesion complex proteins (E-cadherin, vinculin, β-catenin), MDCK cells were lysed with a lysate buffer containing 25 mM NaPIPES, 140 mM NaCl, 1 mM EGTA, 1% CHAPS, 0.5% phosphatidylcholine (PC), 2mM dithiothreitol (DTT) and a cocktail of protease inhibitors (Roche) on ice for 1.5 h. The protein A/G magnetic beads (Cell Signaling Technology) were incubated with either normal rabbit IgG, anti-Piezo1 antibody (1:200) or anti-E-cadherin antibody (1:200) at 4°C for 2 h. Then the antibody-bound beads were incubated with MDCK cell lysate at 4 °C overnight. After incubation, the beads were washed for 5 times with the wash buffer containing 25 mM NaPIPES, 140 mM NaCl, 0.6% CHAPS, 0.14% PC, 2 mM DTT, and finally boiled for 10 minutes in 1 x SDS protein loading buffer. The immune-precipitated proteins were subjected to SDS-PAGE gel and western blotting.

For heterologous overexpression, HEK293T cells grown in a 10 cm-petri dish were transiently transfected with Piezo1-GST, GFP-E-cadherin, β-catenin in a respective amount of 10 μg: 5 μg: 5 μg. For control groups, control vectors were co-transfected to make up the total DNA amount transfected. 40 h after transfection, the confluent cells were lysed using the lysis buffer described above. For GST pull-down experiments, the glutathione magnetic beads (Pierce) were incubated with cell lysates at 4°C for overnight. The beads were washed 3 times with the lysis buffer, then boiled for 10 minutes in 1 x SDS protein loading buffer. The protein samples were subjected to SDS-PAGE and western blotting for analysis.

#### Biotinylation assay

The biotinylation assay was carried out as previously described (Zhang et al., 2017). In brief, HEK293T cells were cultured on poly-D-lysine coated 6-mm dish and transfected with indicated plasmids. After transfection for 48 hours, cells were washed with ice-cold PBS with Ca^2+^ and Mg^2+^ (PBS-CM) for 3 times and incubated with 0.5 mg/mL Sulfo-NHS-LC-Biotin (Thermofisher Scientific) for 45 minutes at 4°C. Then the Sulfo-NHS-LC-Biotin solution was discarded and the reaction was stopped with 100mM glycine solution. Followed by 3 washes with PBS-CM, the cells were lysed with the lysis buffer without DTT. 2% of the lysates were used as the input samples and the remaining cell lysates were incubated with the streptavidin magnetic beads (Thermofisher Scientific) overnight at 4°C. After 5 washes with the wash buffer without DTT, the precipitated sample was boiled for 10 minutes in 1 x SDS protein loading buffer and subjected to SDS-PAGE separation and western blotting.

#### Purification of the Cap fragment protein and interaction with E-cadherin

The purification and characterization of the cap domain (previously termed the C-terminal extracellular domain, CED) was previously described (Ge et al., 2015). The cap fragment protein was induced in Escherichia coli BL21 strain by 0.5 mM isopropyl-β-D-thiogalactoside when the cell density reached an optical density of ~0.8 at 600 nm. After growing at 18°C for 12 h, the cells were collected, washed, resuspended in the buffer containing 25 mM Tris-HCL, pH 8.0, 500 mM NaCl and 20 mM imidazole, and lysed by sonication. The lysates were clarified by centrifugation at 23,000 g for 1 h and the supernatant was collected and loaded onto Ni^2+^ nitrilotriacetate affinity resin (Ni-NTA, Qiagen). The resin was washed extensively with the above buffer and eluted with 280 mM imidazole. The eluate was concentrated and subjected to gel filtration (Superdex-200, GE Healthcare) with the buffer containing 25 mM Tris-HCl, pH 8.0, 200 mM NaCl, 2mM DTT.

For testing the interaction between the purified cap fragment protein and E-cadherin, HEK293T cell were transfected with E-cadherin-Flag construct. 40 h after transfection, the cells were harvested and lysed using the lysis buffer containing 25 mM NaPIPES, 140 mM NaCl, 1 mM EGTA, 1% CHAPS, 0.5% phosphatidylcholine (PC), 2mM dithiothreitol (DTT) and a cocktail of protease inhibitors (Roche) on ice for 1.5 h. The purified cap protein at the desired concentration (4, 8, 16, 32, 64, 128 ng/μl) was added to the aliquots of the cell lysate derived from the E-cadherin-Flag-transfected HEK293T cells. The His-beads were incubated with the cell lysates containing the added cap fragment protein at 4°C overnight, then collected and washed 5 times with the wash buffer. The beads were boiled for 10 minutes in 1 x SDS protein-loading buffer and subjected to SDS-PAGE separation and western blotting.

#### Western blotting

Protein samples from either GST-pull-down or immunoprecipitation were subjected to SDS-PAGE gels for electrophoresis separation. The separated proteins were transferred to 0.45 μm PVDF membranes (Millipore) at 100V for 2h. The membrane was blocked by 5% non-fat milk (Bio-rad) in TBST buffer (TBS buffer with 0.1% Tween-20) for 1 h at room temperature and washed 3 times, 5 min each time with TBST. Then the membrane was incubated with the primary antibodies at 4°C overnight. After 3 times wash with TBST, the membrane was incubated with the peroxidase-conjugated anti-rabbit IgG secondary antibody (CST, 1:10000) or anti-mouse IgG secondary antibody (pierce, 1:10000) at room temperature for 2 h, followed with washing and detection using the enhanced chemiluminescence (ECL) detection kit (Pierce).

#### Immunostaining

Primarily cultured keratinocytes were fixed with 4% PFA for 10 min and then permeabilized with 0.2% Triton for 10 min in room temperature. The cells were blocked with 3% bovine serum albumin (BSA) for 1 h, washed for 3 times, then co-stained with the anti-dsRed antibody (1: 200) and the anti-E-cadherin antibody (1: 200). After washing 5 times, the secondary antibodies Goat anti-Mouse Alexa Fluor-647 (1: 500) and the Donkey anti-Rabbit Alexa Fluor-594 (1:500) were incubated for 2 h at 4°C and washed for 3 times. Phalloidine (100 nM) (Cytoskeleton) was incubated to stain the F-actin fibers. The stained coverslips were mounted onto the glass slide. The imaging procedures were performed on the Nikon A1 confocal microscope with 100 × oil objective (N.A. = 1.49) using the 488, 561, 647 nm excitation wavelength and the 500-550, 570-620, 663-738 nm emission wavelength.

#### RNA interference

Piezo1, E-cadherin and β-catenin knockdown in MDCK cells was achieved by siRNA transfection. MDCK cells were transfected with 50 nM siRNA using Lipofectamine 2000 (Invitrogen) following the manufacturer’s instruction. 48 hours after transfection, the cells were harvested and analyzed with quantitative RT-PCR to detect the knockdown efficiency. The primers for RT-PCR and sequences of siRNA are listed in the key resource table.

#### Membrane-permeable peptides

The peptide of the linker region (amino acids 2171-2185) of the mouse Piezo1 protein was synthesized and myristoylated at its N-terminus (myr-NH2-TEKKYPQPKGQKKKK-COOH) by GeneScript (Nanjing, China) as previously described (Zhang et al., 2017). The scrambled peptide was synthesized with the same composition and did not resemble any known protein (myr-NH2-KQKPKTKEKYKQKGT-COOH). For GST pull-down experiments, the peptides were added to the cell lysates with a working concentration of 200 μM and incubated at 4°C overnight. For the whole-cell patch clamp experiments, the peptides were pre-mixed in the internal solution (200 μM) and filled in the recording pipette.

#### Whole-cell electrophysiology and mechanical stimulation

MDCK cells were briefly digested with trypsin and sparsely re-plated. About 1 h later, the cells were subjected to electrophysiological recordings. HEK293T cells were grown in poly-D-lysine-coated coverslips and transfected with DNA constructs using Lipofectamine 2000 (Thermo Fisher Technology). 36 h after transfection and prior to electrophysiology, the cells were briefly digested with trypsin and sparsely re-plated to obtain individual cells for recording. Electrophysiological recordings normally started about 2 h later after re-plating the cells. For studying the regulatory effect of E-cadherin on wild-type Piezo1 or the mutants, either Flag-E-cadherin-ires-GFP/Piezo1-mRuby or Flag-E-cadherin-mRuby/Piezo1-GST-ires-GFP were co-transfected for identifying cells co-expressing both proteins, which showed both GFP and mRuby fluorescent signals. The observed mechanically activated currents were similar between the two transfection-conditions. For co-transfection, an equal amount of constructs expressing either Piezo1, E-cadherin, or β-catenin were used and a control vector was included to make the total amount of transfected DNA equal among different transfection combinations.

The patch-clamp experiments were carried out with a HEKA EPC10 as previously described (Zhao et al., 2016). For whole-cell patch clamp recordings, the recording electrodes had a resistance of 2–5 MΩ when filled with the internal solution composed of (in mM) 133 CsCl, 1 CaCl_2_, 1 MgCl_2_, 5 EGTA, 10 HEPES (pH 7.3 with CsOH), 4 MgATP and 0.4 Na_2_GTP. The extracellular solution was composed of (in mM) 133 NaCl, 3 KCl, 2.5 CaCl_2_, 1 MgCl_2_, 10 HEPES (pH 7.3 with NaOH) and 10 glucose. All experiments were performed at room temperature. The currents were sampled at 20 kHz, filtered at 2 kHz using Patchmaster software. Leak currents before mechanical stimulations were subtracted off-line from the current traces. Voltages were not corrected for a liquid junction potential (LJP).

Mechanical stimulation was delivered to the cell during the patch-clamp being recorded at an angle of 80° using a fire-polished glass pipette (tip diameter 3–4 μm) as described. Downward movement of the probe towards the cell was driven by a Clampex-controlled piezo-electric crystal micro-stage (E625 LVPZT Controller/Amplifier; Physik Instrument). The probe had a velocity of 1 μm/ms during the downward and upward motion, and the stimulus was maintained for 150 ms. A series of mechanical steps in 1 μm increments was applied every 10 s and currents were recorded at a holding potential of −70 mV. The membrane-permeable linker-peptide and scramble peptide (200 μM) and cytochalasin D (10 μM) were directly added to the recording chamber and incubated for 30 min at 37°C prior to electrophysiological recordings.

#### Cell-attached electrophysiology

Stretch-activated currents were recorded in the standard cell-attached patch-clamp configuration using HEKA EPC10 as previously described (Zhao et al., 2016; Zhao et al., 2018a). The currents were sampled at 20 kHz and filtered at 2 kHz. The recording electrodes had a resistance of 2–3 MΩ when filled with a standard solution consisting of (in mM) 130 NaCl, 5 KCl, 10 HEPES, 1 CaCl_2_, 1 MgCl_2_ and 10 TEA-Cl (pH 7.3,balanced with NaOH). The external solution used to zero the membrane potential consisted of (in mM) 140 KCl, 10 HEPES, 1 MgCl_2_, and 10 glucose (pH 7.3 with KOH). All experiments were carried out at room temperature. The Membrane patches were stimulated with negative-pressure pulses for 500 ms through the recording electrode using a Patchmaster-controlled pressure clamp HSPC-1 device (ALA-scientific). Stretch-activated channels were recorded at a holding potential of −120 mV with pressure steps from 0 to −100 mm Hg (−10 mm Hg increments), and 4 to11 recording traces were averaged per cell for analysis. Current-pressure relationships were fitted with a Boltzmann equation of the form: I(P) = [1 + exp (−(P – P_50_)/s)]−1, where I is the peak of stretch-activated current at a given pressure, P is the applied patch pressure (in mmHg), P_50_ is the pressure value that evokes a half maximal activation of the channel, and s reflects the current sensitivity to pressure.

#### Fura-2 Ca^2+^ imaging

HEK293T cells grown on 8-mm round glass coverslips coated with poly-D-lysine and placed in 48-well plates were transfected with either mRuby-Piezo1 alone or together with Flag-E-cadherin-ires-GFP, and then were subjected to Fura-2 Ca^2+^ imaging 36 h post transfection according to the previously described protocol (Wang et al., 2018). The confluent cells were washed with buffer containing 1 × HBSS (with1.3 mM Ca^2+^) and 10 mM HEPES (pH 7.2 with NaOH), then incubated with 2.5 μM Fura-2-AM (Molecular Probes) and 0.05% Pluronic F-127 (Life technologies) for 30 min at room temperature. After washing with the buffer, the coverslips were mounted into an inverted Nikon-Tie microscopy equipped with a CoolSNAP charge-coupled devise (CCD) camera and Lambda XL light box (Sutter Instrument). GFP positive and -negative cells were selected for measurement of the 340/380 ratio with a 20 × objective (numerical aperture N.A.=0.75) using the MetaFluor Fluorescence Ratio Imaging software (Molecular Device). A stock solution of Yoda1 at 30 mM was solubilized in DMSO and diluted to the desired final concentrations. The compound solutions were perfused into the chamber and cells via a multichannel perfusion system (MPS-2, World Precision Instruments).

For MDCK Ca^2+^ imaging, the cells grown on 8-mm round glass coverslips coated with poly-D-lysine and placed in 48-well plates were transfected with cy3-labeled siRNAs, and then were subjected to Fura-2 single cell Ca^2+^ imaging 36 h post transfection. The confluent cells were washed with buffer containing 1 × HBSS (with1.3 mM Ca^2+^) and 10 mM HEPES (pH 7.2 with NaOH), then incubated with 2.5 μM Fura-2-AM (Molecular Probes) and 0.05% Pluronic F-127 (Life technologies) for 90 min at room temperature. After washing with the buffer, the coverslips were mounted into an inverted Nikon-Tie microscopy equipped with a CoolSNAP charge-coupled devise (CCD) camera and Lambda XL light box (Sutter Instrument). Cy3-labled MDCK cells were selected for measurement of the 340/380 ratio with a 20 × objective (numerical aperture N.A.=0.75) using the MetaFluor Fluorescence Ratio Imaging software (Molecular Device). A stock solution of Yoda1 at 30 mM was solubilized in DMSO and diluted to the desired final concentrations.

#### Reagents

siRNAs specifically targeting canine E-cadherin, Piezo1, β-catenin were synthesized by GenePharma (Nanjing, China). Yoda1 was purchased from pharmacodia. Other chemicals were purchased from Sigma or Ameresco.

#### Quantification and Statistical Analysis

Image processing was carried out by ImageJ. Data in all figures are shown as mean ± S.E.M. Statistical significance was evaluated using either unpaired Student’s t-test or one-way ANOVA for analyzing multiple samples using Prism. Statistical significance was represented as: *p < 0.05, **p < 0.01, ***p < 0.01, ****p < 0.0001. No methods were used to determine whether the data met assumptions of the statistical approach.

#### Data and Code Availability

All relevant data are available from the corresponding author upon reasonable request.

