## Supplementary material for "Tethering Piezo channels to the actin cytoskeleton for mechanogating via the E-cadherin-β-catenin mechanotransduction complex": The key resource table

### KEY RESOURCES TABLE

| REAGENT or RESOURCE | SOURCE | IDENTIFIER |
|---|---|---|
| Antibodies | | |
| mPiezo1 CED antibody | Zhang et al., 2017 | N/A |
| Anti-E-cadherin antibody [M168] | Abcam | Cat # ab76055; RRID: AB_1310159 |
| Vinculin Antibody | Cell Signaling Technology | Cat # 4650; RRID: AB_10559207 |
| β-Catenin Antibody | Cell Signaling Technology | Cat # 9562; RRID: AB_331149 |
| β-Actin (13E5) Rabbit mAb | Cell Signaling Technology | Cat # 4970; RRID: AB_2223172 |
| Anti-GST Antibody | Cell Signaling Technology | Cat # 2622; RRID: AB_331670 |
| Mouse Anti-GFP-Tag Monoclonal Antibody | Abgent | Cat # AM1009a; RRID: AB_352468 |
| Anti-FLAG M2 | Sigma-Aldrich | Cat # F1804; RRID: AB_262044 |
| DsRed Polyclonal Antibody | Clotech | Cat #632496 RRID: AB_10013483 |
| Anti-rabbit IgG, HRP-linked Antibody | Cell Signaling Technology | Cat # 7074; RRID: AB_2099233 |
| Goat anti-Mouse IgG (H+L) Secondary Antibody, HRP | ThermoFisher | Cat # A16066; RRID: AB_2534739 |
| Bacterial and Virus Strains | | |
| Escherichia coli: XL10-Gold Chemical Competent Cell | This paper | N/A |
| Chemicals, Peptides, and Recombinant Proteins | | |
| Dulbecco's Modified Eagle Medium (DMEM) | ThermoFisher | Cat#C11995500BT |
| Fetal Bovine Serum (FBS) | ThermoFisher | Cat#10099141C |
| Penicillin-Streptomycin, Liquid | Gibco | Cat#15140122 |
| polyethylenimines (PEI) | Polysciences | Cat#23966-2 |
| CHAPS | Avanti | Cat#840051C |
| L-α-phosphatidylcholine | Anatrace | Cat#APO129 |
| Protein Inhibitor | Roche | Cat#04693159001 |
| Fura-2 | ThermoFisher | Cat#F1201 |
| Pluronic F-127 | ThermoFisher | Cat# P3000MP |
| Yoda1 | Pharmacodia | N/A |
| Standard Chemical Peptide: P1-2171-2185 | GenScript | N/A |

| | | |
|---|---|---|
| Standard Chemical Peptide: scramble | GenScript | N/A |
| Critical Commercial Assays | | |
| AxyPrep Multisource Total RNA Miniprep Kit 250-prep | Axygen | Cat#AP-MN-MS-RNA-250G |
| RevertAid RT Reverse Transcription Kit | ThermoFisher | Cat#K1691 |
| ClonExpress II One Step Cloning Kit | Vazyme Biotech | Cat#C112-01/02 |
| EZ-Link™ Sulfo-NHS-Biotin | ThermoFisher | Cat#21217 |
| SuperSignal West Pico Chemiluminescent Substrate | ThermoFisher | Cat#34077 |
| Lipofectamine2000 transfection kit | ThermoFisher | Cat#11668019 |
| Experimental Models: Cell Lines | | |
| HEK293T | Xiao et al., 2011 | N/A |
| MDCK | Xu Tan Lab in Tsinghua University | N/A |
| Keratinocytes | Primary Culture | N/A |
| Experimental Models: Organisms/Strains | | |
| Mouse: C57BL/6J | The Jackson Laboratories | Cat# JAX:000664; RRID:IMSR_JAX:000664 |
| Oligonucleotides | | |
| canine Piezo1 RT-qPCR-Forward: GCTCAAGCCCCGAGACATAA | This paper | N/A |
| canine Piezo1 RT-qPCR-Reverse: CACCACCACACCTTCAGTGG | This paper | N/A |
| canine E-cadherin RT-qPCR-Forward: ACACAGGAGTCATCAGCGTG | This paper | N/A |
| canine E-cadherin RT-qPCR-Reverse: GTTAAGCCTTCGCCTTGCAG | This paper | N/A |
| canine β-catenin RT-qPCR-Forward: ACACAGTTCGATGCTGCTCA | This paper | N/A |
| canine β-catenin RT-qPCR-Reverse: ATTGCACGTGTGGCAAGTTC | This paper | N/A |
| canine GAPDH-RT-qPCR-Forward: CCATGTTTGTGATGGGCGTG | This paper | N/A |
| canine GAPDH-RT-qPCR-Reverse: TTGGCTAGAGGAGCCAAGCA | This paper | N/A |
| siRNAs | | |
| Scramble siRNA UUCUCCGAACGUGUCACGU | Genepharma | N/A |

| | | |
|---|---|---|
| canine Piezo1 siRNA GUGCUAUGGUCUCUGGGAUDTDT | Genepharma | Rosenblatt, J. 2017 |
| canine Ecadherin siRNA-871 GCGGAUGAUGAUGUGAAUATT | Genepharma | N/A |
| canine β-catenin siRNA-696 GCACAAUCUUUCUCAUCAUTT | Genepharma | N/A |
| Recombinant DNA | | |
| Plasmid: Piezo1-GST | Ge et al., 2015 | N/A |
| Plasmid: Piezo2-GST | Wang et., 2019 | N/A |
| Plasmid: E-cadherin | Yih Tai Chen and S&M Shasby (Univ. of Iowa) | N/A |
| Plasmid: β-Catenin | Jie Na Lab in Tsinghua | N/A |
| Software and Algorithms | | |
| PatchMaster | HEKA | http://www.heka.com/downloads/downloads_main.html#down_patchmaster |
| GraphPad Prism | GraphPad Software Inc | http://www.graphpad.com/scientific-software/prism/ |
| Origin 9.2 | OriginLab | http://www.originlab.com/ |
| Igor Pro | WaveMetrics | https://www.wavemetrics.com/ |
| Metaflour | Medical Expo | https://pdf.medicalexpo.com/pdf/molecular-devices/metafluor-software-brochure/84611-165738.html |
| The PyMOL Molecular Graphics System | Schrodinger | https://pymol.org/2/ |
